# Quantification of electron transport-related oxidative signals by time and wavelength-resolved redox biosensors and chlorophyll fluorescence

**DOI:** 10.1101/2024.09.09.611861

**Authors:** Matanel Hipsch, Nardy Lampl, Raz Lev, Shilo Rosenwasser

**Author notes:** Corresponding author. (S.R.). These authors contributed equally to this work (Matanel Hipsch, Nardy Lampl, Raz Lev).

## Abstract

Reductive and oxidative signals transmitted from the photosynthetic electron chain to target proteins through the redox signaling network are key regulators of carbon assimilation and downstream metabolism. However, despite their crucial role in activating and inhibiting photosynthetic activity, their relation to photosynthetic efficiency is hardly quantified due to the methodological gap between traditional spectroscopic approaches for investigating photosynthesis and biochemical analyses used in the redox regulation field. Here, we simultaneously quantified redox signals and photosynthetic activity by exploring time and wavelength-resolved fluorescence spectra that capture biosensor and chlorophyll fluorescence signals. Using a set of potato plants expressing genetically encoded redox biosensors, we demonstrated how reductive and oxidative signals are amplified with elevated light intensities and revealed the tight connection between electron transport rate (ETR) and the generation of peroxiredoxin-related oxidative signals. These results demonstrate how full spectrum analysis can pave the way for the integration of genetically encoded biosensors in photosynthesis research and demonstrate light-dependent activation of inhibitory oxidative signals in major crop plants.

## Introduction

The conversion of light into chemical energy through the light and carbon reactions of photosynthesis is a highly complex process that requires continuous coordination between dozens of enzymatic reactions. This coordination ensures adaptation to varying light intensities experienced by plants and avoids the production of harmful ROS. The channeling of reducing and oxidizing equivalents from the photosynthetic electron transport (PET) to regulated thiol proteins plays a significant role in the linkage of the reaction rates of photosynthetic enzymes to instantaneous changes in daytime photon fluxes^1–6^. Reductive signals, catalyzing the reduction of disulfide bonds, are transferred from ferredoxin to Thioredoxins (Trxs) and subsequently redox-regulated target chloroplast proteins via Ferredoxin-Thioredoxin reductase (FTR), or via ferredoxin-NADP(+) reductase (FNR) and NADPH thioredoxin reductase C (NTRC)^7–9^. Counterbalanced oxidative signals, catalyzing disulfide bond formation, are generated by photosynthetically derived hydrogen peroxide (H_2_O_2_) and transmitted through 2-Cys-Prxs, highly efficient thiol peroxidases and high midpoint potential atypical Trxs^10–16^. Accordingly, the redox state of a wide range of photosynthetic enzymes is constantly adjusted through the simultaneous generation of opposing signals, which subsequently dictates the efficiency of light-capturing and carbon-assimilation processes.

Despite the tight link between photosynthetic activity and the redox regulatory network, the relationship between the electron transport rate and the activation of reductive/oxidative signals remains unexplored, mainly due to critical methodological differences in assessing photosynthetic activity versus thiol redox state. While chlorophyll fluorescence and near infra-red absorption measurements are commonly used to non-destructively evaluate the redox state of key photosynthetic components and quantify electron flow through PSII and PSI^17–20^, redox signals transmitted through the redox regulatory network are typically detected using biochemical or proteomics techniques that require protein extracts^21–23^. This methodological gap makes it challenging to directly examine the relationship between the rate of the electron transport and the transmission of signals through the redox signaling network under dynamic environmental conditions.

The development of genetically encoded redox biosensors capable of probing the redox state of key redox components opened new possibilities for studying the relationship between photosynthetic activity and the reductive and oxidative signals^24–27^. Indeed, use of such biosensors has provided key insights into light-dependent changes in chloroplastic GSH redox potential, and in H_2_O_2_, and NADPH levels^28–35^. Moreover, monitoring the real-time redox dynamics of Trx, has been demonstrated during light/dark transition using FRET sensor based on the TRX-targeted chloroplast protein CP12^36^. Similarly, the introduction of chl-roGFP2-PrxΔC_R_, an ultrasensitive chloroplast-targeted 2-Cys peroxiredoxin-based H_2_O_2_ biosensor, based on the yeast roGFP2–Tsa2ΔC_R_ biosensor^37,38^, to Arabidopsis plants, yielded a comprehensive view of inducible photosynthesis-dependent oxidative signals, a phenomenon typically masked by the counterbalanced reductive power^39^.

However, drawing a direct correlation between photosynthetic parameters and biosensor redox state is currently constrained by the standard tools used to analyze biosensor signals, which do not allow recording biosensor and chlorophyll fluorescence while illuminating leaves with actinic light over various light intensities. Accordingly, integrating redox biosensors into photosynthesis research requires fluorescence detection methods that are comparable with spectroscopic approaches used for investigating photosynthesis.

In this work, the Arabidopsis chl-roGFP2-Prx and chl-roGFP2-PrxΔC_R_ biosensors were introduced into potato plants to explore the relationship between light intensity, photosynthetic parameters, and the generation of photosynthetically derived oxidative signals. A newly developed system for time-resolved and wavelength-resolved fluorescence measurements^40^ was exploited to map chlorophyll fluorescence-derived photosynthetic parameters and biosensor oxidation state under the exact same lighting setup and at an equivalent resolution. The presented results provide insights into the relationship between electron transport rates and the generation of reductive and oxidative signals over a wide range of light intensities and illustrate how biosensor-derived data can be integrated into photosynthesis research by capturing full fluorescence spectra.

## Results and Discussion

### Generation of potato plants expressing chl-roGFP2-Prx and chl-roGFP2-PrxΔC_R_

chl-roGFP2-PrxΔC_R_ is a biosensor generated by the genetic fusion of roGFP2 to a mutated version of Arabidopsis 2-Cys Prx A (BAS1), in which the resolving cysteine (C_R_) was changed to alanine^39^. Integration of the Arabidopsis 2-Cys peroxiredoxin A signal peptide targets the biosensor to the chloroplast stroma^41^. Comparison of the light responsiveness of chl-roGFP2-PrxΔC_R_, chl-roGFP2-Prx (without the mutation in the C_R_) and the unfused chl-roGFP2 showed that the absence of TRX-dependent reduction of chl-roGFP2-PrxΔC_R_ allowed for quantitative, real-time readout of photosynthetically-derived oxidation signals^39^

Potato plants were chosen as a model to measure photosynthetically derived oxidative signals in agriculturally significant plant systems due to their global importance for food security^42^ and the availability of highly efficient transformation protocols. To monitor oxidizing signals, agrobacterium-mediated genetic transformation was performed to obtain potato plants cv Desiree expressing chl-roGFP2-PrxΔC_R_ and chl-roGFP2-Prx (Fig. 1a). Efficient chloroplast targeting was verified by the overlap of the chl-roGFP2-PrxΔC_R,_ chl-roGFP2-Prx and chlorophyll fluorescence signals (Fig. 1b) as was also shown for potato plants expressing chl-roGFP2 (Fig. 1b^34^). The overall fluorescence intensity recorded for roGFP2-PrxΔC_R_ and chl-roGFP2-Prx was lower than chl-roGFP2 (Fig, 1c), a phenomenon typically observed in plant lines expressing fused protein biosensors. No phenotypic differences between independent lines expressing the biosensors and wild type were observed; lines with the highest fluorescence intensity were selected for subsequent experiments.

**Figure 1:**
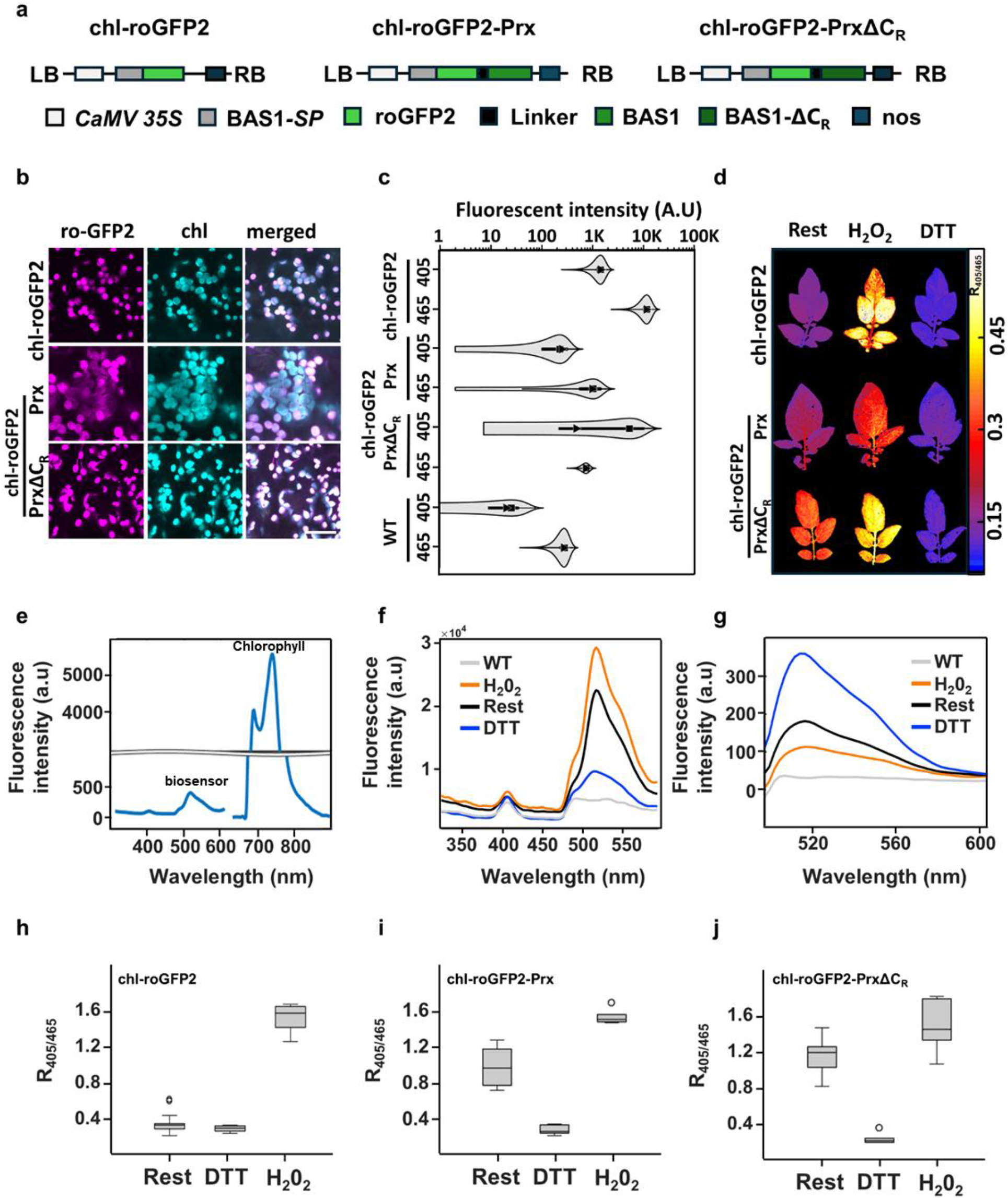
Generation of potato plants expressing chl-roGFP2-Prx and chl-roGFP2-PrxΔC_R_, and fluorescence detection using live imaging and full-spectrum acquisition. (a) Schematic diagram of the gene cassette used to introduce chl-roGFP2, chl-roGFP2-Prx and chl-roGFP2-PrxΔC_R_ into potato plants. (b) Confocal images showing the colocalization of fluorescence signals emanating from potato plants expressing the biosensors and chlorophyll fluorescence. (chl-roGFP2 excitation: 488 nm). (c) Pixel intensity distributions for chl-roGFP2 and chl-roGFP2-Prx, chl-roGFP2-PrxΔCR and WT. The data are displayed as a violin plot, showing mean pixel intensity from 3-4 leaves per line. The plot includes the range within 1.5 times the interquartile range (IQR), with a box plot inside indicating the 25th to 75th percentiles. The ◼ marks the mean, and the ▴ marks the median (d) Ratiometric analysis conducted by dividing pixel by pixel fluorescence images taken following excitation at 405 and 465 nm during rest and under fully oxidized (1000 mM H_2_O_2_, 10 min) and fully reduced [100 mm dithiothreitol (DTT), 60 min] conditions. Images of detached leaves were digitally combined for comparison. (e) Simultaneous detection of biosensor and chlorophyll fluorescence by wavelength-resolved acquisition. Spectra derived from chl-roGFP2-PrxΔC_R_ expressing plants is shown. The presented spectrum was derived from the data of two spectrometers tuned at different sensitivity levels. (f, g) Spectral analysis of chl-roGFP2-PrxΔC_R_ plants during rest, under fully oxidized and fully reduced conditions following excitation at 405nm (f) and 465nm (g). (h-j) Fluorescence ratios (405/465) in plants expressing chl-roGFP2, chl-roGFP2-Prx, chl-roGFP2-PrxΔC_R_ during rest, under fully oxidized and fully reduced conditions following excitation. Biosensor-derived fluorescence signals were calculated by integrating fluorescence intensities over the 500-550nm range.

In vivo fluorescence ratiometric imaging of leaves treated with H_2_O_2_, dithiothreitol (DTT) or untreated (rest) was conducted to examine the sensitivity of chl-roGFP2-PrxΔC_R_ and chl-roGFP2-Prx toward redox changes. As shown in Fig. 1d, images generated by dividing pixel-by-pixel images acquired following 405-nm and 465-nm excitations (R_405/465_) demonstrated higher R_405/465_ values in both plant lines in leaves treated with H_2_O_2_ compared with DTT. Notably, while rest R_405/465_ was close to DTT in chl-roGFP2, showing a highly reduced state of the probe, higher oxidation states of chl-roGFP2-PrxΔC_R_ and chl-roGFP2-Prx under rest were evident (Fig. 1d). These results, which demonstrate the direct oxidative effect of 2-Cys Prx on the roGFP moiety, are consistent with probe characteristics previously observed in yeast and *Chlamydomonas reinhardtii* expressing roGFP2–Tsa2ΔC_R_^37,38^ and in Arabidopsis plants expressing chl-roGFP2-PrxΔC_R_ and chl-roGFP2-Prx^39^.

### Wavelength-resolved measurements capture biosensors and chlorophyll fluorescence

Parallel recording of the redox state of the three biosensors under dynamic light conditions was expected to constitute a powerful tool for both assessing the activity of 2-Cys Prx-dependent oxidative signals and deducing the reductive activity of NTRC/TRXs and GSH/Grx pathways. Redox alterations in response to light were measured using a newly developed system that allows time-resolved and wavelength-resolved fluorescence measurement while applying red actinic light at varying intensities, including a burst of single turnover saturating flashes (STF) applied to “close” PSII reaction centers^40^. As shown in Fig 1e, spectrally resolved data facilitated the capture of fluorescence emission at 500-550 nm, i.e., signals emanating from chl-roGFP2-PrxΔC_R_ plants excited at 405 nm, as expected for eGFP-based biosensors. Notably, resolving the entire spectra allowed for the simultaneous capture of biosensor and chlorophyll fluorescence, enabling the correlation between the biosensor oxidation state and chlorophyll-fluorescence-driven photosynthetic parameters.

Resolving the entire spectra also enabled the assessment of the structured autofluorescence originating from plant pigments. The emission signal peaked at 490 nm in wild-type (WT) plants excited at 405 nm (Fig. 1f and Supp. Fig.1). This signal was used to quantitively assess the structural autofluorescence bleed-through into the biosensor channels by calculating the factor between the autofluorescence measured at 490 nm and the bleed-through to the roGFP emission channels. Subsequently, roGFP-associated fluorescence values were corrected by subtracting the computed autofluorescence, which improved the accuracy and reliability of subsequent analyses (See Methods).

Shifting chl-roGFP2-PrxΔC_R_ to its fully oxidized state by applying H_2_O_2_ caused an increase in emission (calculated as the area under the curve between 500 nm and 550 nm) after excitation at 405 nm and decreased when excited at 465 nm, compared to untreated plants (Fig. 1 f&g). On the other hand, leaves treated with DTT to shift the biosensor to its fully reduced state showed a less intense signal upon 405 nm excitation and increased brightness upon 465 nm excitation (Fig. 1 f&g). The dynamic range (R_405/465_ for fully oxidized state divided by R4_05/465_ for fully reduced state) for each sensor was calculated after conducting similar experiments in plants with chl-roGFP2-Prx and chl-roGFP biosensors (Fig.1 h-j). Dynamic ranges of 5-6 were recorded for chl-roGFP2-Prx, chl-roGFP2-PrxΔCR, and chl-roGFP2, consistent with the performance of the biosensors in Arabidopsis plant^34,39^.

### Biosensor oxidation patterns reflect the combined activity of reducing and oxidizing signals

To analyze the dynamics of reducing and oxidative signals at various light intensities, dark-adapted biosensor-expressing plants were exposed to a series of incremental light intensities with a 2-fold change between two consecutive light steps (0, 20, 40,80,160, 320, and 640 µmol m^2^s^−1^), covering the low to high light intensity range^43^. Each illumination step lasted for 10 min to allow for short-term acclimation. Full spectra recording was used to measure biosensor signals and chlorophyll fluorescence-derived parameters under the exact conditions (Fig. 2a-c). To ensure minimal interference with photosynthetic activity during the fluorescence measurements, integration time for biosensor and chlorophyll fluorescence were set to 15-10,0.7 ms, respectively. Saturating pulses were applied to measure PSII operating efficiency (ΦPSII), non-photochemical quenching (NPQ), and Electron transport rate (ETR). To capture the changes occurring immediately after light intensity modifications, the frequency of measurements was highest immediately after light intensity was shifted and was gradually decreased as the acclimation period progressed (Supp. Fig. 2).

**Figure 2:**
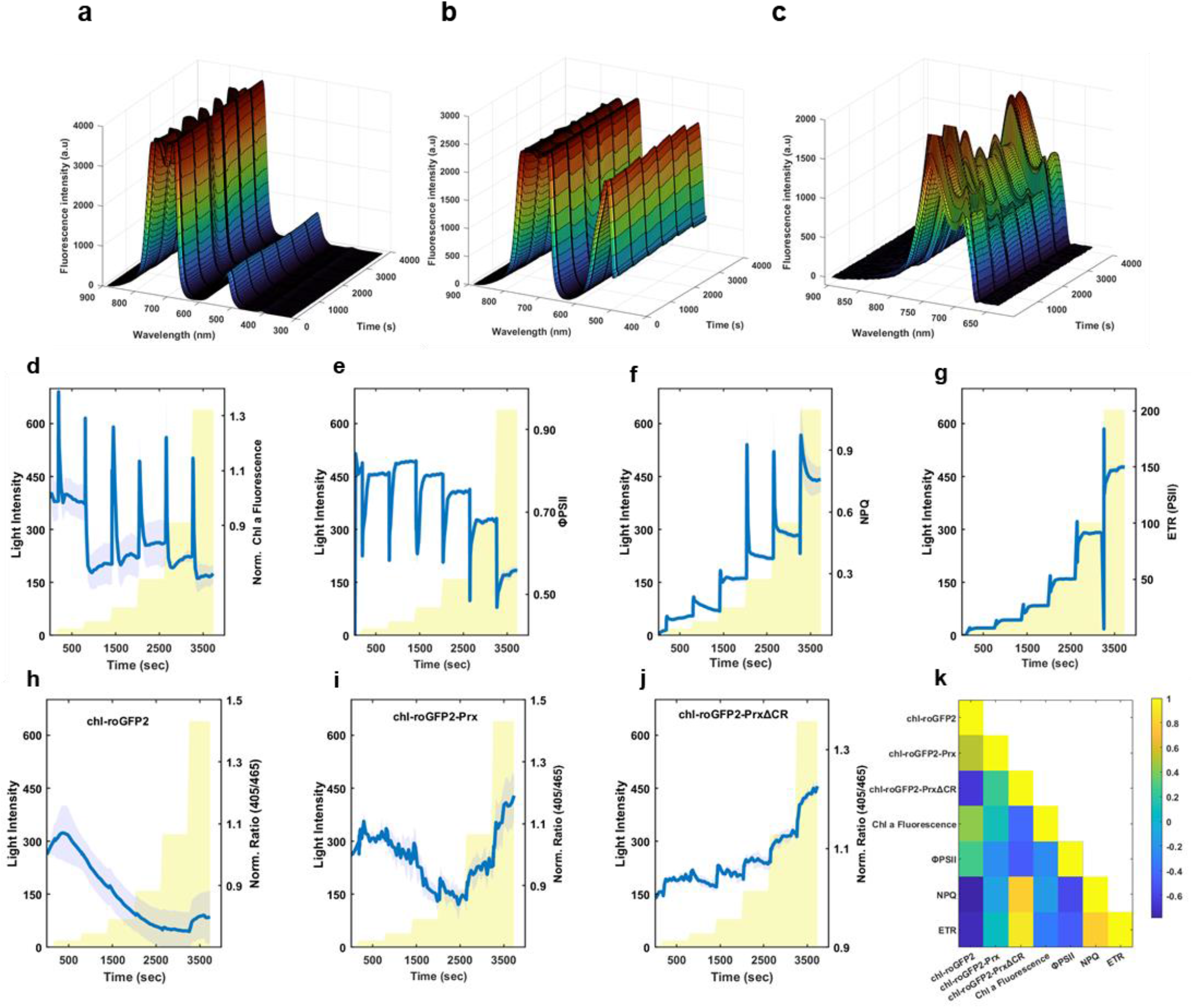
Spectra capturing biosensor and chlorophyll fluorescence in response to increases in actinic light intensity. (a-c) 3D plots displaying time-dependent spectra of chl-roGFP2 plants excited at 405 nm (a), 465 nm (b) for biosensor detection, and 600 nm (c) for chlorophyll fluorescence. Signals of biosensors (a, b) and chlorophyll fluorescence (c) were detected with two different spectrometers optimized for signal intensities. (d-g) Chlorophyll fluorescence-derived parameters, including normalized fluorescence (d), ΦPSII (e), NPQ (f) and ETR II (g) extracted from spectra analysis and following the application of saturating pulses. Plants were exposed to 2 fold increases in light intensities ranging from 0−640 µmol m^2^s^−1^. Each illumination step lasted for 10 min. Light intensity is indicated in the yellow bars. Values represent means ± SE obtained from at least three independent experiments. (h-j) Biosensors ratio (405/465) values extracted from spectra, in response to light intensity increments. Biosensor names are shown on the graphs. Biosensor-derived fluorescence signals were calculated by integrating fluorescence intensities over the 500-550nm range. Plants were exposed to the same light conditions as in d-g, as indicated by the yellow bars. Values represent means ± SE obtained from at least three independent experiments. (k) Liner relationships between electron transport and biosensor signals. An analysis of linear correlation coefficients (r) between photosynthetic parameters derived from chlorophyll fluorescence and biosensors oxidation state is presented.

As shown in Fig. 2d, increasing light intensities induced rapid chlorophyll fluorescence peaks, reflecting the reduction of the Qa site, followed by adaptation of the photosynthetic machinery to the newly experienced light intensity either by induction of Qa oxidation or activation of non-photochemical quenching. The similar amplitudes observed at each light transition are consistent with the recently observed fold-change response of Chl a fluorescence to light^43^. Decreasing PSII values ranging from 0.81 to 0.56, at 20 µmol m^2^s^−1^ and 640 µmol m^2^s^−1^, respectively, accompanied by increasing NPQ values ranging from 0.08 to 0.75, were recorded (Fig. 2e&f). As expected for light intensities below light saturation, increasing in ETR II, ranging from 6.8 to 150at 20 µmol m^2^s^−1^ and 640 µmol m^2^s^−1^, respectively, was also documented (Fig. 2g).

The overall response of chl-roGFP2 and chl-roGFP2-Prx to the increasing light intensities up to 160 µmol m^2^s^−1^ was similar and involved a light-dependent reduction in the redox state. Higher light intensities induced chl-roGFP2-Prx oxidation, while no change occurred in chl-roGFP2 in response to 320 µmol m^2^s^−1^, followed by a slight increase in the last light step of 640 µmol m^2^s^−1^ (Fig. 2h&i). No such pattern was observed in plants exposed to the same sequence of measurements but without illumination (Supp. Fig. 3), emphasizing the light-dependency of the observed patterns. The redox state of redox biosensors is shaped by the counterbalanced activity of reductive and oxidative signals emanating from the photosynthetic electron transport chain. Accordingly, the different turning points toward oxidation observed in chl-roGFP2-Prx compared to chl-roGFP2 reflect the prevalence of Prx-associated oxidative signals at 320 µmol m^2^s^−1^, which attenuate the reduction activity of TRXs and GSH/GRXs. These results reflect the predominant action of reductive signals in response to incremental light intensities in low to medium light intensity range and the higher activity of oxidative signals at higher light intensities.

A contrasting oxidation dynamic was observed for the chl-roGFP2-PrxΔC_R_, which underwent oxidation as light intensities increased, showing a step increase in the generation of 2 Cys Prx-related oxidative signals at light intensities above 80 µmol m^2^s^−1^ (Fig. 3j). The observed oxidation was likely attenuated by the reductive activity of GSH/GRXs, as observed in the chl-roGFP2 line (Fig. 2h). Accordingly, the chl-roGFP2-PrxΔC_R_ was normalized to the chl-roGFP2 (Supp. Fig. 4, Oxidation Index [OI]), refining the oxidative activity of 2 Cys Prx. An increase in OI was clearly observed in response to the incremental rise in light intensities in the low to medium light range, with no response in the last step (from 320 to 640 µmol m^2^s^−1^). These results demonstrate the generation of Prx-related oxidative signals at habitual light intensities and the oxidation of GSH pool at moderate to high intensities. In light of the inhibitory effects of Prx-related oxidative signals on Calvin-Benson cycle enzymes, these results highlight the limitations of photosynthesis imposed by oxidative signals in one of the world’s most important crops.

Electron flow to photosystem I (PSI) was shown to be the prominent source of H_2_O_2_ in chloroplasts, with the acceptor-side capacity of PSI suggested to regulate the rate of superoxide formation through the Mehler reaction^44,45^. Notably, fully reduced PSI acceptors and overwhelming of the antioxidant pathways of the water-water cycle have been suggested to play a role in chloroplast-derived ROS signaling resulting from H_2_O_2_ accumulation^45^. Nonetheless, the quantifiable oxidative signals under light levels below stressful levels suggest that while H_2_O_2_ export from the chloroplast only occurs under stress conditions associated with PSI damage, oxidative signals transmitted to target proteins through 2 Cys Prx are delivered ‘below the radar’ of the antioxidant systems. Accordingly, the parallel induction of chl a fluorescence and chl-roGFP2-PrxΔC_R_ oxidation indicate the high sensitivity of 2 Cys Prx to changes in the electron transport rate, suggesting a molecular mechanism for adaption of photosynthetic reaction rates to short-term and transitory light changes.

A correlation matrix was constructed to explore the relationships between biosensor signals and photosynthetic activity. The tight association between the electron transport rate and generation of oxidative signals was further demonstrated by the strong correlation between ETR and chl-roGFP2-PrxΔC_R_ oxidation state (r= 0.91, Fig. 2k). A significant negative correlation (r= -0.73) was also found between chl-roGFP2 and ETR. These results demonstrate that reductive and oxidative signals are tightly connected to electron transfer. A positive correlation was also found between ETR II and ETR I and the reductive state of two key Calvin-Benson-Cycle (CBC) enzymes, FBPase and SBPase^46^. Consequently, either the activation of CBC enzymes by reduction or their inhibition by oxidation is initiated by the electron transport chain, with increasing light intensities amplifying both signals.

While it is possible that the dominant reductive activity largely attenuates the oxidative signals at a steady state, differences in the kinetics of both signals can introduce a new layer of regulation during the light transition, underlining the importance of oxidative signals. This context-dependent modulation can be harnessed to achieve precise responses to changes in light intensities. In addition, differential sensitivity of target proteins to both signals, possibly mediated by specific Trxs, may create a continuum of protein redox states, ensuring fine-tuned control and specificity within the redox signaling system.

Taken together, parallel quantification of chlorophyll and genetically-encoded redox biosensors fluorescence using spectrally resolved emission data demonstrate the strong correlation between reductive and oxidative signals and electron transport rates. Given the importance of redox signaling in dictating photosynthetic efficiency, integration between classical spectroscopic photosynthetic measurements and genetically encoded biosensors is likely to provide a novel perspective on how light reactions are regulated under dynamic environmental conditions.

## Methods

### Plant material, growth conditions and experimental setup

Wild type (Solanum tuberosum) and potato plants expressing chl-roGFP2, chl-roGFP2-Prx, and chl-roGFP2-PrxΔC_R_ were planted in moist soil and vegetatively propagated from cuttings in a controlled-environment greenhouse. Several days before experiments, 3-4-week-old plants were moved to a FytoScope FS-RI 1600 plant growth chamber (Photon Systems Instruments). Plants were cultivated in the chamber under light conditions of 300 µmol m^2^s^−1^, with ambient CO2 levels and relative humidity (R.H.) maintained at 60-70%. Plants were dark-adapted from 2 h before measurements.

### Production of transgenic plants

Binary vectors containing the chl-roGFP2-Prx, chl-roGFP2-PrxΔC_R_ genes^39^ were used for Agrobacterium-mediated infection of potato leaves (cv. Désirée). Briefly, chemically synthesized genes, including the Arabidopsis 2-Cys Prx A (AT3G11630) signal peptide, roGFP2 and Arabidopsis 2-Cys Prx A were cloned into the binary vector pART27, using the NotI restriction enzyme and transformed into GV3101 Agrobacterium tumefaciens^39^.

For potato transformation, leaves were taken from a clean culture and infected by Agrobacterium tumefaciens strain LBA 4404 harboring the pART27 plasmids, which contain the chl-roGFP2-Prx, chl-roGFP2-PrxΔC_R_ constructs^34,47,48^. After inoculation, cultures were transferred to A cali induction medium containing kanamycin 50 mg/L, cefotaxime 500 mg/L, 6-benzylaminopurine 0.1 mg/L and 1-naphthaleneacetic acid 5 mg/L, for 10 days. Then, plants were transferred to shot induction medium (regeneration medium) containing zeatin-riboside (Z.R.) 2 mg/L, NAA 0.02 mg/L, gibberellic acid 3 0.02 mg/L and appropriate antibiotics. The medium was refreshed every 14 days until shoots appeared. Fully regenerated transgenic explants, lacking roots, were transferred from cultures to moist soil for rooting and further sorting. Appropriate lines were selected by evaluating the chl-roGFP2 fluorescence signal.

### Confocal microscopy

Confocal microscopy images were acquired with a Leica DMI4000 confocal system (Leica Microsystems) and the LAS X Life Science Software, using a HCX APO U-V-I 40x/0.75 DRY UV objective. Images were acquired at a 1024 4096×1024 4096-pixel resolution, with a 507-534 nm emission bandpass and PMT gain of -830 628.2(V) following excitation at 488nm or 405nm for roGFP2 fluorescence detection. A 652-692nm emission bandpass and PMT gain of -571.4(V) following excitation at 488nm were used to image chlorophyll fluorescence. Merged images were generated using LAS X software.

### Whole-plant imaging

Whole-plant fluorescence was detected using an Advanced Molecular Imager H.T. (Spectral Ami-HT, Spectral Instruments Imaging, LLC., USA), and images were acquired using the Aura software. For chl-roGFP2, chl-roGFP2-Prx and chl-roGFP2-PrxΔC_R_ fluorescence detection, excitation was performed with 405 nm±10 or 465 nm±10 LED light sources and a 515 nm±10 emission filter was used. For chlorophyll detection, a 405 nm±10 LED light source and 670 nm±10 emission filter were used. All images were captured under the following settings: exposure time = 1s, pixel binning =2, field of view (FOV) = 15cm, LED excitation power 40% and 60%, for 405 nm, 465 nm excitations, respectively. Excitation power for chlorophyll detection was 5%. Chlorophyll autofluorescence was measured to generate a chlorophyll mask, which was then used to select pixels that returned a positive chlorophyll fluorescence signal. Only those pixels were subsequently considered for the roGFP analysis. For background correction, autofluorescence measured from WT plants was used to assess the factor between the autofluorescence measured at 448 nm and the bleed-through to the roGFP emission channels^49^. This factor was used to generate a scaled version of the autofluorescence, which was subtracted from the original roGFP images. Ratiometric images were created by dividing, pixel-by-pixel, the corrected 405 nm image by the corrected 465 nm image and displaying the result in false colors. Images were pre-processed using a custom-written MATLAB script.

### Full-spectrum analysis

Time-dependent biosensor and chlorophyll fluorescence signals were captured with a ChloroSpec spectrometer (ChloroSpec B.V., The Netherlands)^40^, using the ChloroSoft software. ChloroSpec was equipped with 3 LED sources at 405/20nm, 465/20 nm, and 650/20 nm and two spectrometers that cover a spectral range of 313 nm to 885 nm. In order to maximize the detection of biosensor signals and avoid saturation of chlorophyll fluorescence, the two spectrometers were set at different sensitivity levels. Short-pass filters of 460nm, 475nm, and 660nm were placed in front of the 405, 465, and 650 LEDs, respectively, to prevent the excitation light from entering the spectrometer. The red LED was used for actinic light and saturating pulses. For ch-roGFP2, the 405 nm and 465 nm excitations were delivered for 10 ms, at an intensity of 2000 μmol m^−2^ s^−1^, while chl-roGFP2-Prx, and chl-roGFP2-PrxΔC_R_ were excited with a 15 ms pulse at an intensity of 3500 μmol m^−2^ s^−1^. Data were analyzed with the ChloroKin Software and a custom-written MATLAB script. Biosensor emissions were calculated by integrating the signal between 500 nm and 550 nm. For background correction, autofluorescence measured from WT plants was used to assess the factor between the autofluorescence measured at 490 nm and the bleed-through to the roGFP emission channels (500-550nm). This factor was used to subtract autofluorescence values from biosensor-derived signals. Reaction center closing was achieved by applying a burst of STF (250 pulses, each 130 μs wide, separated by 3 ms intervals) of 60,000 μmol m^−2^ s^−1^ red excitation light. ΦPSII and NPQ values were calculated based on chlorophyll fluorescence intensity detected by photodiode at 685nm. Changes in total chlorophyll fluorescence signals were detected using multi-turnover flash (MTF) pulses of 6,000 μmol m^−2^ s^−1^ for 0.7ms. Chlorophyll fluorescence and biosensor signals were recorded in two different experiments to avoid the possible effect of saturating pulses on biosensor readout.

## Supporting information

Supplementary Images 1-4

## Acknowledgments

We would like to thank Prof. Alfred R. Holzwarth for integrating the biosensor module into Chloropsec, as well as for his invaluable assistance throughout our work with the system. This research was supported by the European Research Council (ERC-COG, AGRIREDOX, grant no. 101086608) and the Israel Science Foundation (grant No. 1779/21).

## Author contributions

M.H., N.L., R.L. and S.R. performed the research and analyzed the data. S.R. wrote the manuscript with all authors providing valuable feedback and comments on the draft.

## Notes

### Competing Interest Statement

The authors have declared no competing interest.

