## Supplementary Images 1-4 for "Quantification of electron transport-related oxidative signals by time and wavelength-resolved redox biosensors and chlorophyll fluorescence"

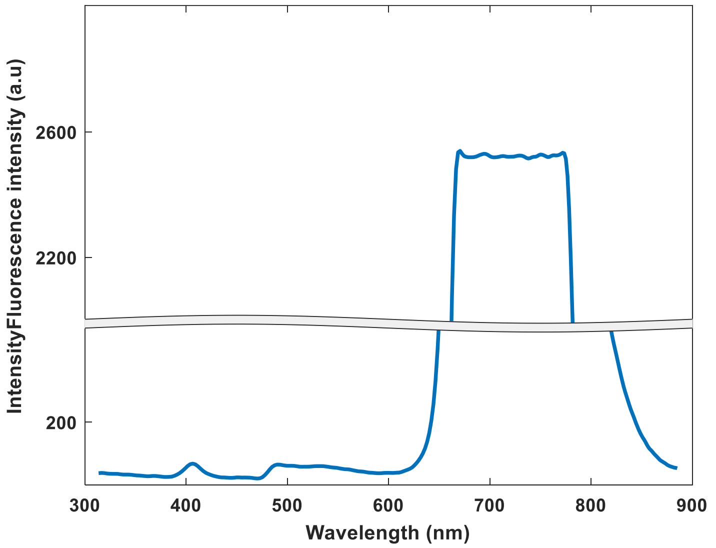


**Supp. Fig.1**: Spectra derived from chl-roGFP2-PrxΔCR expressing plants excited at 405nm is shown. The observed peak at 490nm was used to assess autofluorescence emanated from plant pigments (See Methods).

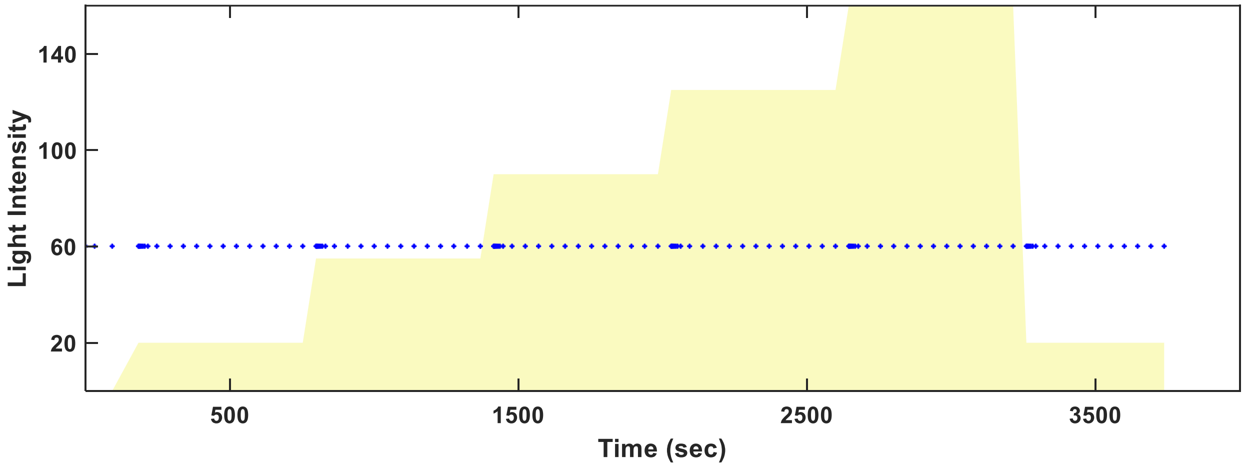


**Supp. Fig.2**: **Light conditions and measurement frequency applied in Fig.2**. Each blue dot represents a single measurement. As shown, the measurement frequency was highest immediately following a change in light conditions.

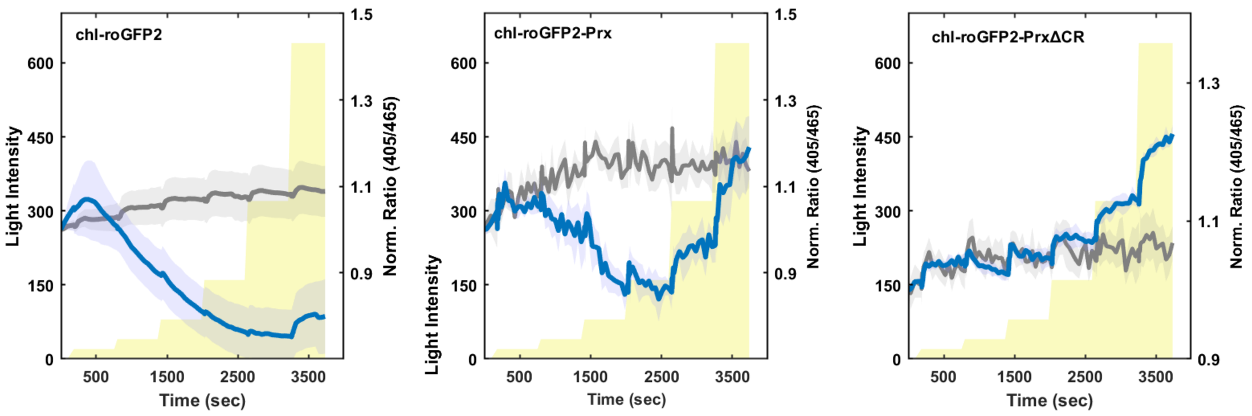


**Supp. Fig.3**: **Constant dark control.** Biosensor ratio (405/465) values extracted from spectra, in response to light intensity or constant dark increments are shown. The blue line represents plants exposed to light increments, and the gray line represents plants exposed to constant darkness.

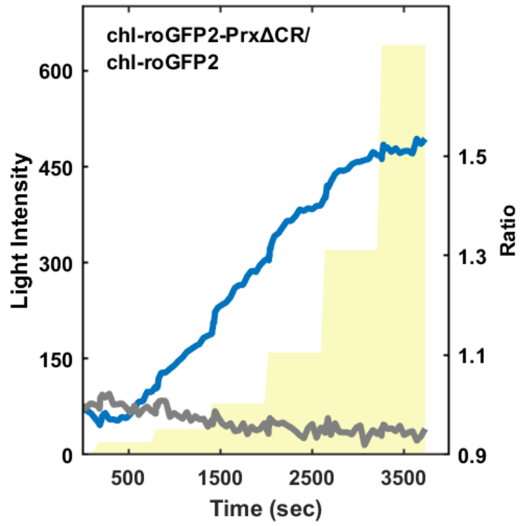


**Supp. Fig.4: Changes in Oxidation index in response to light intensity increments.** The Oxidation index was calculated by normalizing chl-roGFP2-PrxΔCR to the chl-roGFP2 data on Fig. 2 h&j. The blue and gray lines represent values calculated from plants exposed to light increments or constant darkness, respectively.
